# Blood-brain barrier disrupting stimuli induce production of extracellular vesicles with distinct protein cargoes and functionality

**DOI:** 10.64898/2026.01.15.699203

**Authors:** Alec Z. Xiang, Zara Adeel, Andrew J. Shih, Betty Diamond

## Abstract

The blood-brain barrier (BBB) protects the brain from substances in the circulation; yet, this barrier can be breached. Systemic lupus erythematosus is a disease in which over 50% of patients experience manifestations of neuropsychiatric lupus (NPSLE), with many displaying BBB dysfunction. High levels of TNFα, IL-1β, C5a, and epinephrine, which have all been shown to cause BBB permeability, may be present in the serum of SLE patients.

Brain microvascular endothelial cells (BMVECs) are a major structural element of the BBB. We asked whether BMVECs stimulated with barrier-disrupting factors release extracellular vesicles (EVs) which might participate in autocrine signaling or might have utility as diagnostic biomarkers of BBB permeability. We analyzed proteins in EVs secreted by unstimulated and stimulated BMVECs by mass spectrometry and used secreted EVs for further BMVEC stimulation. We subjected BMVECs to bulk RNA-seq to identify response signatures to the initial stimulus and to secondary stimulation with EVs. We show that there are agent-specific EV-associated protein profiles. EVs from BMVECs subjected to BBB-breaching stimuli alter the BMVEC transcriptional program, representing a potential feed-forward mechanism. Finally, we suggest that the proteins associated with EVs might be markers of BBB disruption.

## Introduction

The blood-brain barrier (BBB) protects the brain from neurotoxic substances in the circulation. There are, however, agents that can cause impairments in BBB integrity. In systemic lupus erythematosus (SLE), an autoimmune disease characterized by production of autoantibodies including brain-reactive antibodies, and systemic inflammation, central nervous system (CNS) manifestations such as cognitive dysfunction and mood disorder are common.

Disruption of the BBB with subsequent exposure of brain parenchyma to neurotoxic antibodies and other soluble factors within the circulation is thought to be one mechanism for development of these neuropsychiatric manifestations of disease^1^.

The BBB normally maintains a careful homeostasis between the CNS and the peripheral circulation, regulating small molecule and ion concentrations while excluding most cells and proteins. It consists primarily of brain microvascular endothelial cells (BMVECs), which form the major structural and functional component of the brain’s vasculature. Astrocytes and pericytes are also important components of the BBB, and act in concert to regulate cerebral blood flow and vascular tone while also communicating with BMVECs to confer BBB-specific properties to these cells^2,3^. The consequences of BBB disruption may be drastic, and can include toxic activation of astrocytes, vascular pathology, microglial activation, and chronic pathological changes in the neurons of the CNS^4,5^. A BBB breach in SLE can result from high circulating levels of several pro-inflammatory molecules such as TNFα and IL-1β, from C5a, a complement component generated through activation of the complement cascade, as well as from epinephrine.

BMVECs, like other cells, secrete EVs which may be altered in response to molecules in the circulation. They can secrete EVs either into the systemic circulation or the CNS parenchyma, depending on whether the vesicles originate from the apical or basolateral aspect^6^. The secreted vesicles may have potential utility as biomarkers of BBB dysfunction. EVs from BMVECs and endothelial progenitor cells already have been shown to exacerbate BBB dysfunction in inflammatory contexts, but may also participate in maintenance and repair of the BBB^6,7^.

We compare the protein cargoes of EVs secreted in response to BBB-breaching stimuli, describe the transcriptional profile of the BMVECs secreting the EVs, and explore the potential of these EVs for autocrine signaling through characterization of transcriptional responses to EVs secreted by stimulated BMVECs. To work toward a circulating EV-based diagnostic of BBB health and function, we sought to determine whether EVs secreted by activated BMVECs contained both stimulus-specific proteins and proteins common to EVs following all stimulations.

## 2 Materials and Methods

### 2.1 Brain Microvascular Endothelial Cell Culture and Stimulation

The hCMEC/D3 cell line was purchased from Sigma-Aldrich (Sigma-Aldrich, St. Louis, MO, SCC066) and thawed according to the manufacturer’s instructions. Cells were cultured in 12-13 mL of Microvascular Endothelial Cell Growth Medium-2 (EBM-2MV) from Lonza (Lonza, Walkersville, MD 00190860) supplemented with 3% heat-inactivated complete fetal bovine serum (FBS; R&D Systems, Minneapolis, MN, S11150H), 1.4 µM hydrocortisone (Sigma-Aldrich, St. Louis, MO, H0135-1MG), 1 ng/mL human basic fibroblast growth factor (bFGF/hFGF-B) (Sigma-Aldrich, F9786-5ML), 5 µg/mL ascorbic acid (Sigma-Aldrich, A4403-100MG), 1% penicillin/streptomycin, 10mM HEPES (Thermo Fisher, Waltham, MA 15630-080), and 1% chemically defined lipid concentrate (Thermo Fisher, 11905031).

For primary stimulations, serum-free medium was prepared using Gibco Human Endothelial SFM (Thermo Fisher, 11111044) supplemented with 1.4 µM hydrocortisone (Sigma-Aldrich, H0135-1MG), 1 ng/mL human basic fibroblast growth factor (bFGF/hFGF-B) (Sigma-Aldrich, F9786-5ML), 5 µg/mL ascorbic acid (Sigma-Aldrich, A4403-100MG), 1% penicillin/streptomycin, 10mM HEPES (Thermo Fisher, 15630-080), and 1% chemically defined lipid concentrate (Thermo Fisher, 11905031). Medium was refreshed every 3 days.

Cells were grown to 75-90% confluence prior to stimulation, and all cells were counted and viability assessed by trypan blue staining. TNFα (Peprotech, Cranbury, NJ 300-01A), IL-1β (Peprotech, 200-01B), and C5a (Peprotech, AF-300-70) were diluted to 1 mg/mL in sterile ultrapure H_2_O, aliquoted, stored at -80°C, and thawed immediately before use. Cells were either cultured in serum-free medium alone or stimulated for 12 hours with TNFα at 25 ng/mL, IL-1β at 10 ng/mL, and C5a at 50 ng/mL. Epinephrine bitartrate (Sigma-Aldrich, E4375-1G) was diluted to 200 nM and likewise used for 12-hour stimulations. All primary stimulations were performed in T75 flasks.

For secondary stimulations, BMVECs were exposed to EVs secreted by the BMVECs in primary stimulations. Secondary stimulations were performed in supplemented serum-free medium. Cells were grown to 75-90% confluence in T25 flasks and treated with 1*10^9^ EVs per flask (4*10^7^ EVs/cm^2^) resuspended in 4 mL of supplemented serum-free medium for 12 hours.

### 2.2 Blood-Brain Barrier Permeability Assessment

To create a bilayer BBB model, hCMEC/D3 immortalized human BMVECs were co-cultured with primary human astrocytes, a generous gift from the laboratory of Dr. Joan W. Berman (Albert Einstein College of Medicine, Bronx, NY, USA) on 0.2% gelatin-coated Corning 3.0µm transwell inserts as described previously (Eugenin and Berman, 2003)^8^. Briefly, endothelial cells were grown in the upper chamber of Transwell^®^ inserts, which were inverted and coated with primary astrocytes from culture passage 4-5. Several inserts in each batch were tested to ensure formation of an impermeable BBB through addition of Evans Blue-conjugated albumin (EBA), which was prepared according to previously described methods (Eugenin and Berman, 2006)^9^. Coated inserts were cultured in phenol red-free DMEM without FBS and with 0.45% EBA at 200 µL per upper chamber, with the addition of TNFα at 25 ng/mL, IL-1β at 10 ng/mL, C5a at 50 ng/mL, epinephrine bitartrate (Sigma-Aldrich, E4375-1G) at 200 nM, 0.5mM EDTA (Fisher Scientific, 17892) as a positive control, or simply EBA-containing medium alone. The lower chamber contained 400 μl of phenol red-free DMEM/10% FBS. After 3h of incubation at 37°C, medium in the lower chamber was collected and its fluorescence intensity at emission 620 nm /excitation 680 nm was measured to quantify the passage of EBA through the BBB model.

### 2.3 Extracellular Vesicle Enrichment

Extracellular vesicles were enriched from cell culture supernatant through successive centrifugation and ultracentrifugation. An initial centrifugation at 300g for 5 minutes was performed to pellet cells. Supernatant was retained and centrifuged at 2000g for 10 minutes, followed by transfer of supernatant to a clean tube and centrifugation at 3200g for 30 minutes to pellet cellular debris and organelles.

Supernatants were then ultracentrifuged in a Sorvall WX 80+ ultracentrifuge (Thermo Fisher, 75000080) for 100,000g for 2 hours to pellet small EVs. Supernatant was decanted and droplets were gently aspirated prior to resuspending the pellet by pipetting 40 times and rinsing with 0.1µm-filtered PBS and centrifuging for 100,000g for an additional 1.5 hours. Pellets were resuspended in 0.02µm-filtered PBS by vigorously pipetting 60 times, while taking care to avoid bubbles.

Biofluids from which EVs were obtained were either processed immediately or stored at - 80°C. Stored samples were thawed on ice prior to EV enrichment. Freeze-thaw cycles were minimized, with EVs subjected to no greater than two (2) freeze-thaw cycles.

### 2.4 ZetaView NTA

Extracellular vesicle size distributions were acquired on a ZetaView TWIN NTA (Particle Metrix, Inning am Ammersee, Germany PMX-220), calibrated as per manufacturer’s instructions using 100nm polystyrene beads (Alpha Nano Tech, Durham, NC 23A1009) diluted in MilliQ water at 1:250,000. Daily performance assessments were also run prior to data acquisition to ensure that trueness and precision are reported as “Very Good” prior to initiating sample analysis. Samples were diluted at a range of 1:10 – 1:1000 in MilliQ water immediately prior to loading into the sample chamber. For acquisition of size distributions, concentrations, and zeta potentials, the camera was set to a sensitivity of 80, shutter speed of 100, and framerate of 30 frames per second. Chamber temperature was set at 22.0°C. We maintained optimal focus throughout acquisition and rinsed the chamber between each sample with at least 5mL of MilliQ water, regularly inspecting to ensure that no bubbles were present. Inclusion and exclusion of any data was performed according to automatic selection criteria of the instrument.

### 2.5 Transmission Electron Microscopy

EVs were fixed to a glow-discharged 400 mesh copper grid (Electron Microscopy Sciences, Hatfield, PA) for two minutes and negatively stained with 1% uranyl acetate prior to being observed in a FEI Tecnai 20 microscope (Hillsboro, OR) at 120kV. Images were captured on a TVIPS TemCam-F416 4kx4k camera (TVIPS GmbH, Gauting, Bayern) with EM-Menu 4 software for image collection.

### 2.6 Western Blot

Prior to running gels, µBCA or standard BCA kits (Thermo Fisher) were used to quantify protein concentrations for equilibration of protein content between lanes. Pre-cast Novex 4-12% Bis-Tris mini-gels (Thermo Fisher) were loaded with equivalent amounts of protein and run at 120V for 2h prior to being transferred to nitrocellulose for 2h at 30V.

The following antibodies from BD Biosciences (Franklin Lakes, NJ), Abcam (Waltham, MA), and Thermo Fisher were diluted in TBS-T and used for Western blots: Flotillin-1 at a dilution of 1:500 (BD Biosciences, 610820), CD81 at a dilution of 1:500 (Abcam, ab79559), and donkey anti-mouse HRP secondary antibody at a dilution of 1:3500 (Thermo Fisher, A16011).

Membranes were developed using an Advansta ECL kit (San Jose, CA) and imaged on an Amersham ImageQuant 800 (Cytiva, Marlborough, MA) with images captured every thirty seconds over the course of five minutes.

### 2.7 Mass Spectrometry

All EVs from stimulations with TNFα, IL-1β, C5a, and epinephrine were submitted as a single batch. Suspended EVs were dried and re-dissolved in 80uL 8M Urea/20mM DTT/50mM ammonium bicarbonate at RT for 45min. Free cysteines were alkylated with iodoacetamide (45min at RT in the dark). Urea was diluted by adding 150uL 50mM ammonium bicarbonate followed by 5ug LysC in 20uL. Samples were allowed to digest o/n. In the morning, samples were further diluted by adding 200uL 50mM ammonium bicarbonate and a 2nd digestion with trypsin (1ug trypsin in 20uL) was carried out for 6h. Digestion was halted by addition of neat trifluoroacetic acid. Peptides were solid-phase extracted prior to analysis by direct-loading nano-LCMS. 80 min analytical gradient and separated using a 25cm/75um EasySprayer column (C18 reversed phase). Mass spectrometer (Ascend operated in high res./high mass accuracy mode). The data were processed using MaxQuant v 2.4.2.0 and searched against a Human database (2024) concatenated with common contaminants. Matched peptides were filtered using a false discovery rate of 1% or better and it was required that a protein was measured in 3-of-3 replicates in at least one condition. Further analysis was performed using the Limma package in R.

The data from EV mass spectrometry are derived from the quantification of proteins contained within EVs isolated from the supernatant of BMVECs which have undergone primary stimulation as described in 4.1. The mass spectrometry proteomics data have been deposited to the ProteomeXchange Consortium via the PRIDE^10,11^ partner repository with the dataset identifier PXD072771.

### 2.8 RNA Isolation

Cells were processed as per manufacturers’ instructions using the Qiagen RNeasy Mini Kit (Qiagen, Germantown, MD 74106). In summary, cells were lysed in Qiagen RLT buffer (Qiagen 79216) and stored at -80°C if not immediately processed. Lysates were homogenized using the QIAshredder kit (Qiagen 79656) prior to RNA isolation, including treatment with DNAse, on RNeasy columns. RNA was eluted into Eppendorf LoBind tubes (Eppendorf, Enfield, CT 022431102) in RNAse-free water, and RNA concentration and OD260/280 were measured using the NanoDrop spectrophotometer (Wilmington, DE). All samples with OD260/280 ≥2.0 were submitted to Novogene (Sacramento, CA) for sequencing using the Illumina (San Diego, CA) NovaSeq platform with a paired-end 150 bp sequencing strategy as a single batch.

### 2.9 Bulk RNA Sequencing and Analysis

Raw fastq files corresponding to paired-end reads for each sample were downloaded from Novogene and their integrity verified with an MD5 checksum. Quality control was performed using FastQC and reads were aligned using STAR aligner to the hg38 reference genome downloaded from Ensembl (database version 112.38/GENCODE 46), along with the corresponding GTF file (version 112)^12,13^. Ensembl IDs were converted to gene symbols using biomaRt and any ID without a gene symbol was removed^14,15^. Subsequent differential expression analysis was performed using R statistical software (v4.4.1; R Core Team 2024) with the DESeq2 package and log fold changes shrunk using “ashr”^16,17^. Heatmaps were plotted using the R package pheatmap and Venn diagrams using ggvenn. The data discussed in this publication have been deposited in NCBI’s Gene Expression Omnibus (GEO) and are accessible through GEO Series accession number GSE315626 (https://www.ncbi.nlm.nih.gov/geo/query/acc.cgi?acc= GSE315626).

### 2.10 Statistics

Values depicted on heatmaps are standard deviations of gene expression for each condition from the mean scaled gene expression scores derived from controls. Multiple comparisons were made by ANOVA with appropriate post-tests. Unpaired two group/sample data (non-parametric) was analyzed by t-test. P values of 0.05 or less were considered significant.

Genes and proteins with an absolute value of log 2-fold change (log2FC) compared to control > 0.5 and adjusted p-value < 0.05 were considered differentially expressed genes (DEGs) and adjusted p-value < 0.05 considered differentially expressed proteins (DEPs).

## 3 Results

### 3.1 Analysis of EV-Associated Proteins

To validate that we could analyze intact EVs, we first isolated EVs from cell culture supernatant and visualized them by transmission electron microscopy (TEM) to ensure that they exhibited the classical morphological finding of a cup-shaped disk, providing evidence of the presence of phospholipid bilayers, (Figure S1A) and EV markers Flotillin-1 and CD81 on Western blot (Figures S1B-D). We then determined whether BMVECs exposed to BBB-breaching stimuli would have alterations in the number, size, or contents of the EVs they secreted.

To verify whether BMVECs were functionally altered by the various stimuli which have been described to cause a BBB breach, specifically TNFα, IL-1β, C5a, and epinephrine, we exposed co-cultures of hCMEC/D3 BMVECs and primary human astrocytes to these stimuli (Figure 1A). We found that barrier integrity measured by permeability to EBA was decreased at 3 hours, confirming that these agents act as effective disruptors of the BBB. All stimuli at the concentrations used throughout subsequent experiments breached the BBB in this coculture system to an extent comparable to the positive control, 0.5mM EDTA, while culture medium with EBA alone did not induce a breach. In BMVEC monocultures, stimulation with TNFα led to a 1.71-fold increase in EV number per cell compared to unstimulated cells (Figure 1B). No increase was observed with any other stimulation condition. Mean particle diameter remained consistent among all stimulation conditions (Figure 1C).

**Figure 1.**
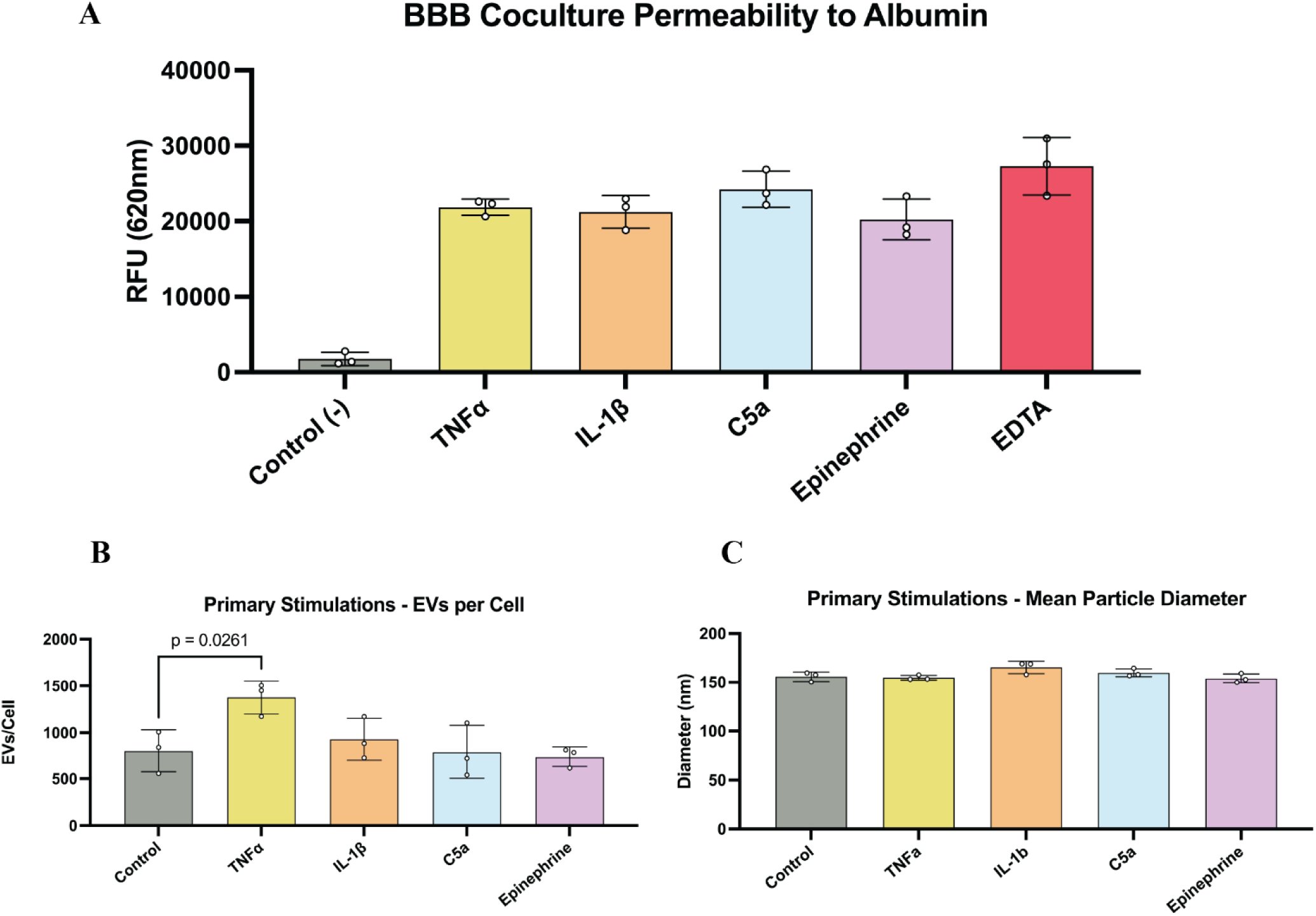
Activation of BMVECs with stimulants. A) Permeability of hCMEC/D3 cells and primary astrocyte cocultures to Evans Blue-labeled albumin following 3h of stimulation with the indicated stimulations at the same concentrations used in hCMEC/D3 monocultures. B) Number of EVs isolated from cell culture supernatant following 12h of indicated stimulations normalized against cell number. Stimulation with TNFα resulted in a significant increase in EV production (p=0.026). Other stimulations did not yield significant changes in EV secretion per cell. C) Median particle size is consistent among stimulations. Each dot represents data from one T75 flask of stimulated hCMEC/D3 cells, with mean ± SD shown for each group. Three flasks were analyzed for each stimulation.

EVs secreted by BMVECs exposed to TNFα, IL-1β, C5a, and epinephrine were collected from the supernatant and their protein content was analyzed by mass spectrometry, revealing 2319 unique differentially expressed proteins (DEPs). A PCA of the protein expression data for each stimulation condition was plotted (Figure 2A). TNFα stimulations cluster on one extreme of the first principal component accounting for 17.8% of variance. IL-1β and C5a cluster together on one end of PC2, accounting for 15.5% of variance. We generated a heatmap for those proteins which were unique to each stimulation condition while also being within the top 100 differentially expressed proteins for that condition (Figure 2B). There were 64 EV-associated proteins which met these criteria in TNFα stimulations, 12 in IL-1β stimulations, 7 in C5a stimulations, and 20 in epinephrine stimulations. We plotted the overlap between the EV-associated proteins secreted by BMVECs of each stimulation condition and those secreted by unstimulated control BMVECs (Figure 2C-F) and found that between 84.8-86.8% of all proteins were common between control EVs and stimulation-derived EVs.

**Figure 2.**
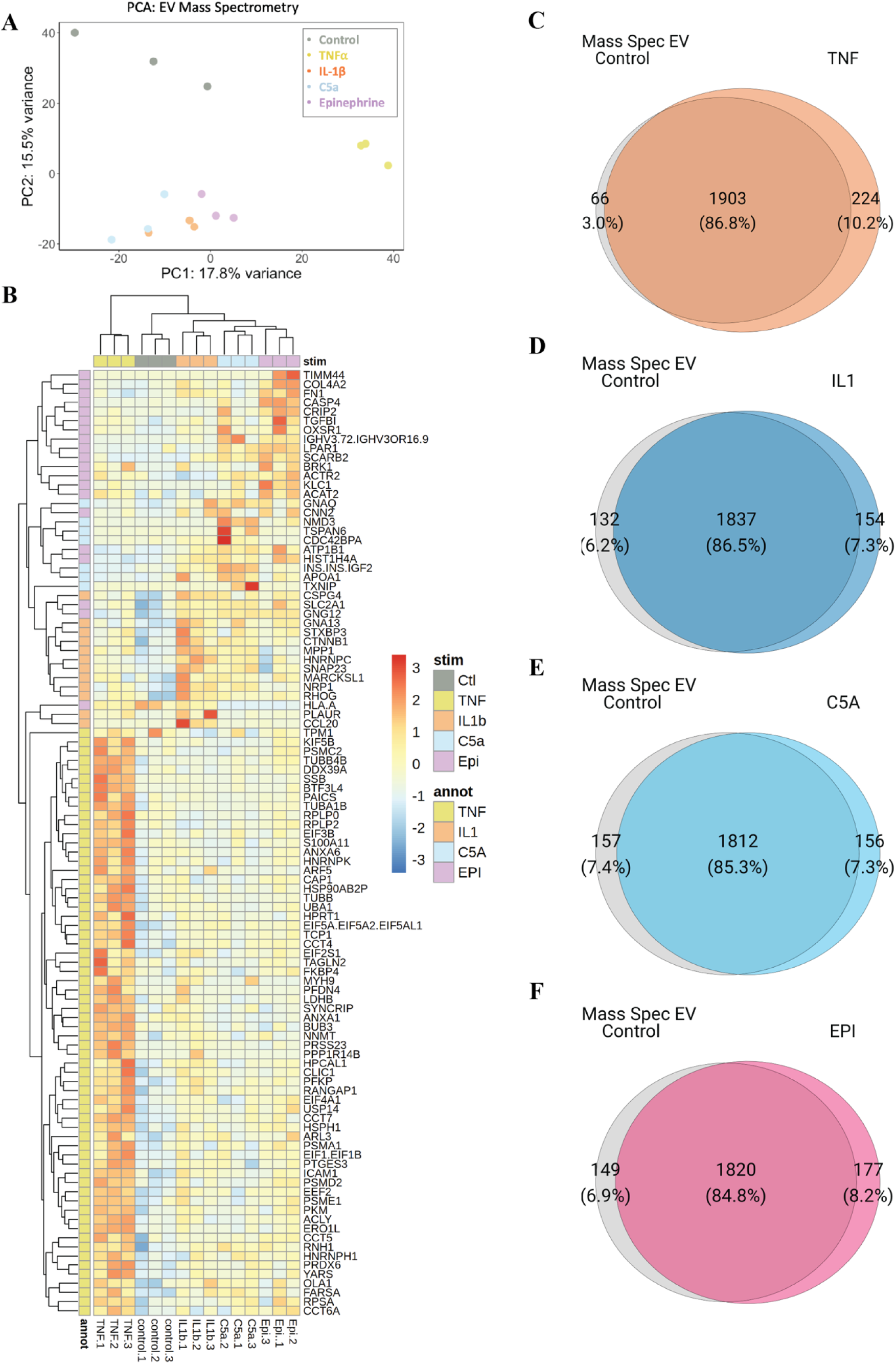
BMVEC EV-associated proteins vary with different stimuli. A) PCA plot of EV-associated proteins from BMVECs which were unstimulated or exposed to the indicated stimulus. The proportion of variance represented by each principal component is noted in parentheses. B) Heatmap depicting EV-associated proteins which are both within the top 100 differentially expressed proteins and also unique to each stimulation condition. C-F) Venn diagrams depicting, from left to right in each diagram, the number of proteins detected only in EVs from unstimulated cells; the number of proteins which were detected in both control EVs and stimulation EVs; and the number of proteins which were detected only in the stimulation conditions C) TNFα, D) IL-1β, E) C5a, and F) epinephrine

EVs secreted by TNFα and IL-1β stimulated cells contained several interferon (IFN) signaling-related proteins, namely, STAT1, UAP1, CNP, RANGAP1, CCL20, and the IFN-inducible protein MX1. We also identified several EV-associated proteins which were common to all stimulation conditions: CCT2, CCT3, PPA1, UAP1, MARCKS, HN1L, TMSB10, EPB41L3, NRAS, EPB41L2, and FERMT2. The proteins CCT2, CCT3, PPA1, UAP1, MARCKS, and HN1L were within the top 100 DEPs of each stimulation.

### 3.2 Transcriptional Profiles of Primary Stimulations

Following culture without stimulation with TNFα, IL-1β, C5a, or epinephrine for 12 hours, BMVECs were harvested and their RNA extracted for bulk RNA-seq. Principal component analysis revealed the formation of several distinct clusters, with PC1 accounting for 49% of variance and PC2 accounting for 13% (Figure 3A). Controls sat at the lower bound of PC1 and the upper bound of PC2. TNFα-stimulated samples were at the upper bound of PC1, while IL-1β stimulated samples were at the lower bound of PC2. Triplicates correlated strongly with one another.

**Figure 3.**
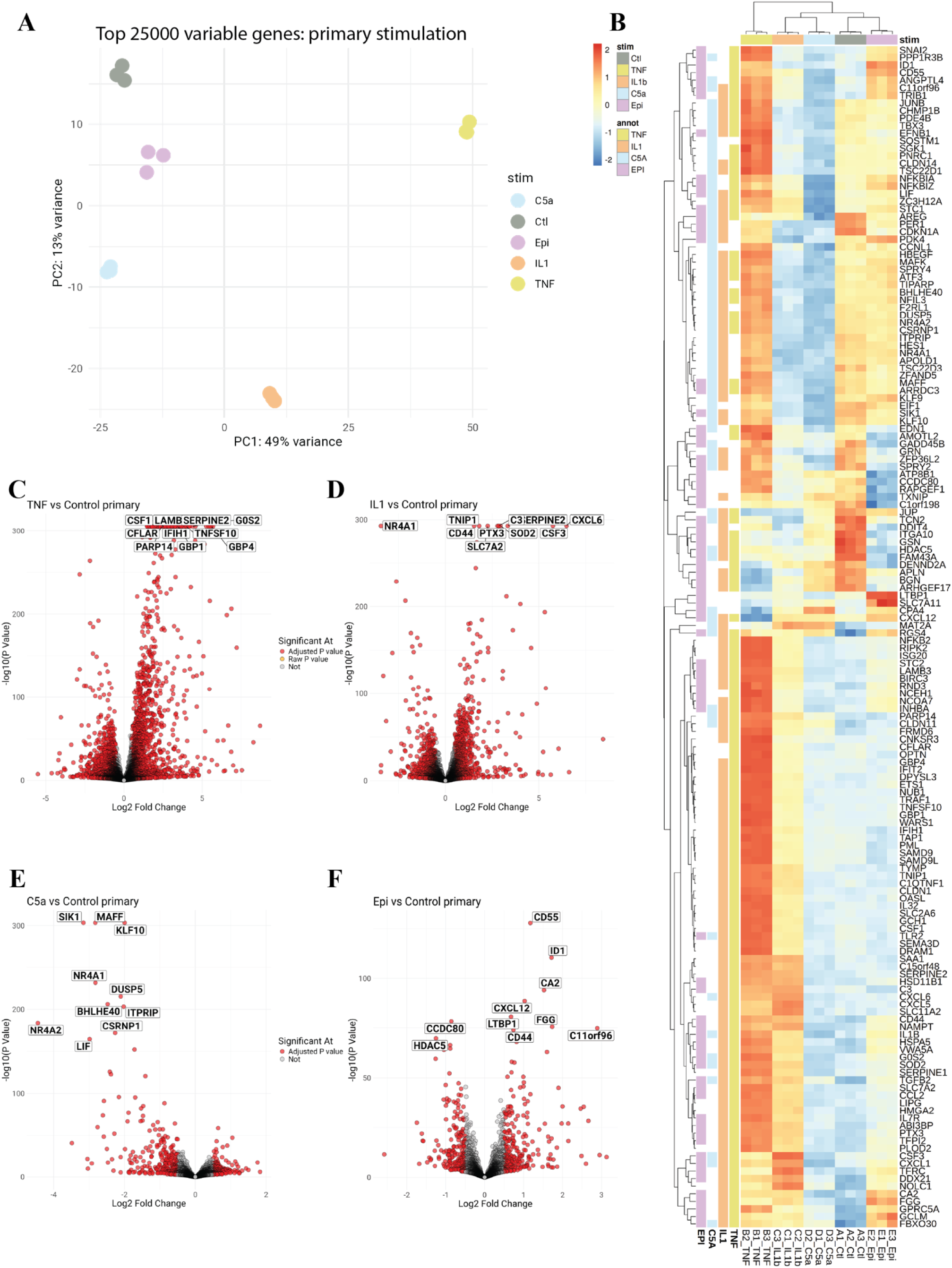
Primary stimulations induced stimulus-dependent differential gene expression. A) PCA plot generated with the top 25000 most variable genes among all primary stimulations (n=3 for each condition). B) Heatmap depicting the top differentially expressed genes among all stimulations. Sample annotations are colored by their stimulation and gene annotations are colored if they are significant at all in that stimulation. C-F) Volcano plots of differentially expressed genes of primary stimulations for C) TNFα, D) IL-1β, E) C5a, and F) epinephrine compared to unstimulated cells. The top 10 differentially expressed genes, sorted by adjusted P value, are labeled.

DEGs in each stimulation were identified and plotted on a heatmap (Figure 3B). Distinct clusters of strongly downregulated genes are notable in C5a and IL-1β stimulations. We identified seventeen DEGs expressed with the same directionality across all stimulation conditions and within the top 100 DEGs of at least one condition (Table S1). Components of the pro-angiogenic EGFL signaling pathway, *AREG* and *EGFL7,* were downregulated across all conditions. Some of the differentially regulated genes are likely critical in barrier disruption, such as the tight junction protein *CLDN5* which was downregulated across all conditions, and the matrix metalloproteinase (MMP) *MMP2* which was upregulated across all stimulations. Volcano plots reveal that the most differentially expressed genes of each stimulus are unique in their identities (Figure 3C-F), consistent with the corresponding heatmap. Transcriptomic profiles for primary stimulations with TNFα and IL-1β were more distinct than the protein profiles of the stimulated EVs (Figure 4).

**Figure 4.**
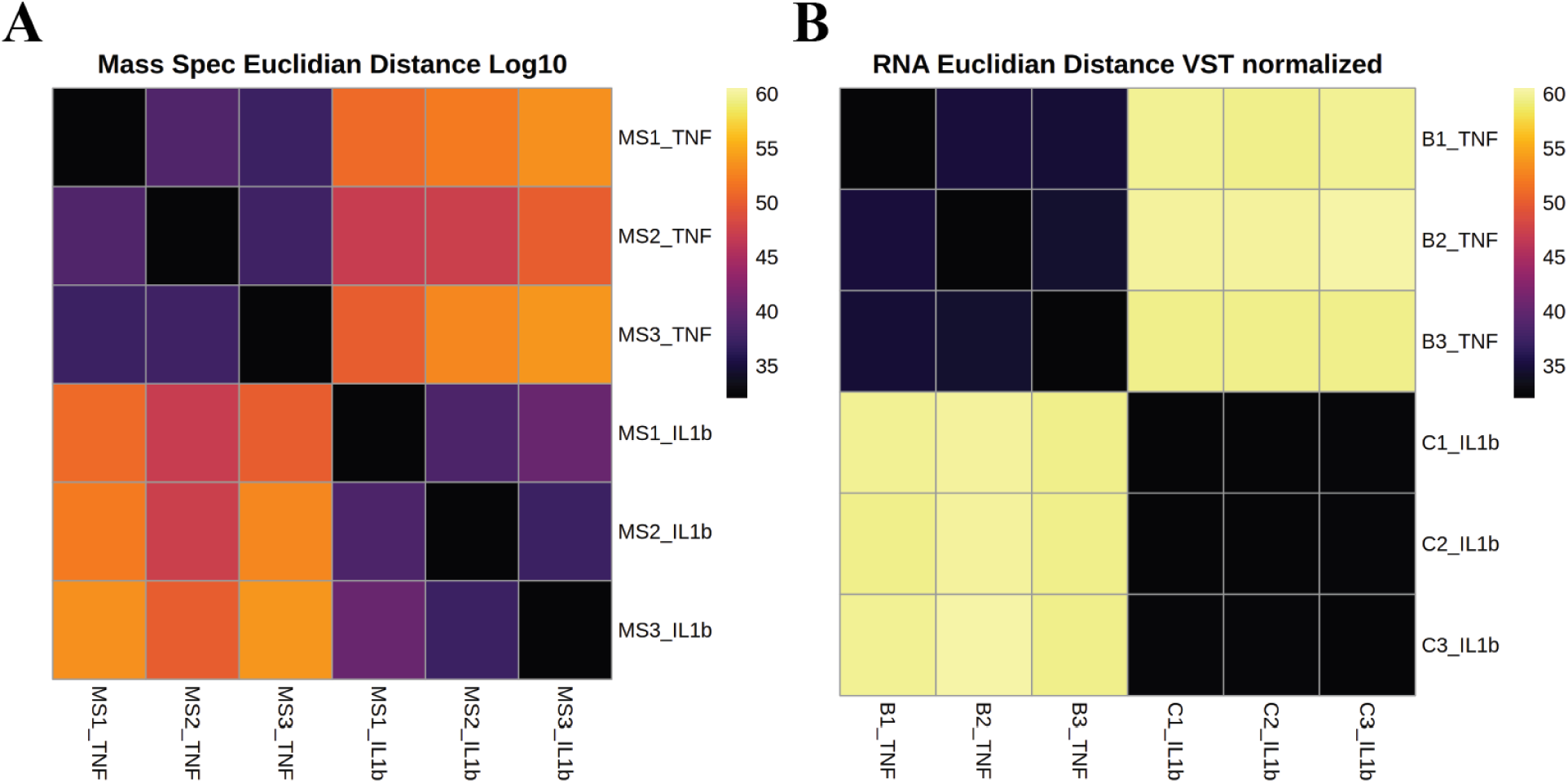
Correlations between transcripts and EV proteins in TNFα and IL-1β stimulations. A) Euclidean distance plot of EV proteins from TNFα and IL-1β primary stimulations. B) Normalized Euclidean distance plot of transcripts from TNFα and IL-1β primary stimulations. Darker colors for both plots indicate a higher similarity between samples while a lighter color indicated less similarity.

### 3.3 Secondary Stimulations

To assess whether EVs secreted by activated BMVECs are capable of independently acting as modulators of BMVECs, we treated BMVECs with equivalent numbers of EVs following primary stimulations. Transcriptional profiles of these secondary stimulations revealed three distinct clusters on a PCA plot.

TNFα and IL-1β formed one cluster with C5a and controls formed another, with the epinephrine samples now exhibiting a distinct pattern of gene expression (Figure 5A). gene expression profiles among these secondary stimulations were more similar than the profiles following primary stimulations, albeit with a lower number of DEGs (Figure 5B). Notably, the identities and directionality of several of the most differentially expressed genes are the same in TNFα and IL-1β secondary stimulations, although the magnitude of gene expression differs between these two conditions (Figure 5C, D).

**Figure 5.**
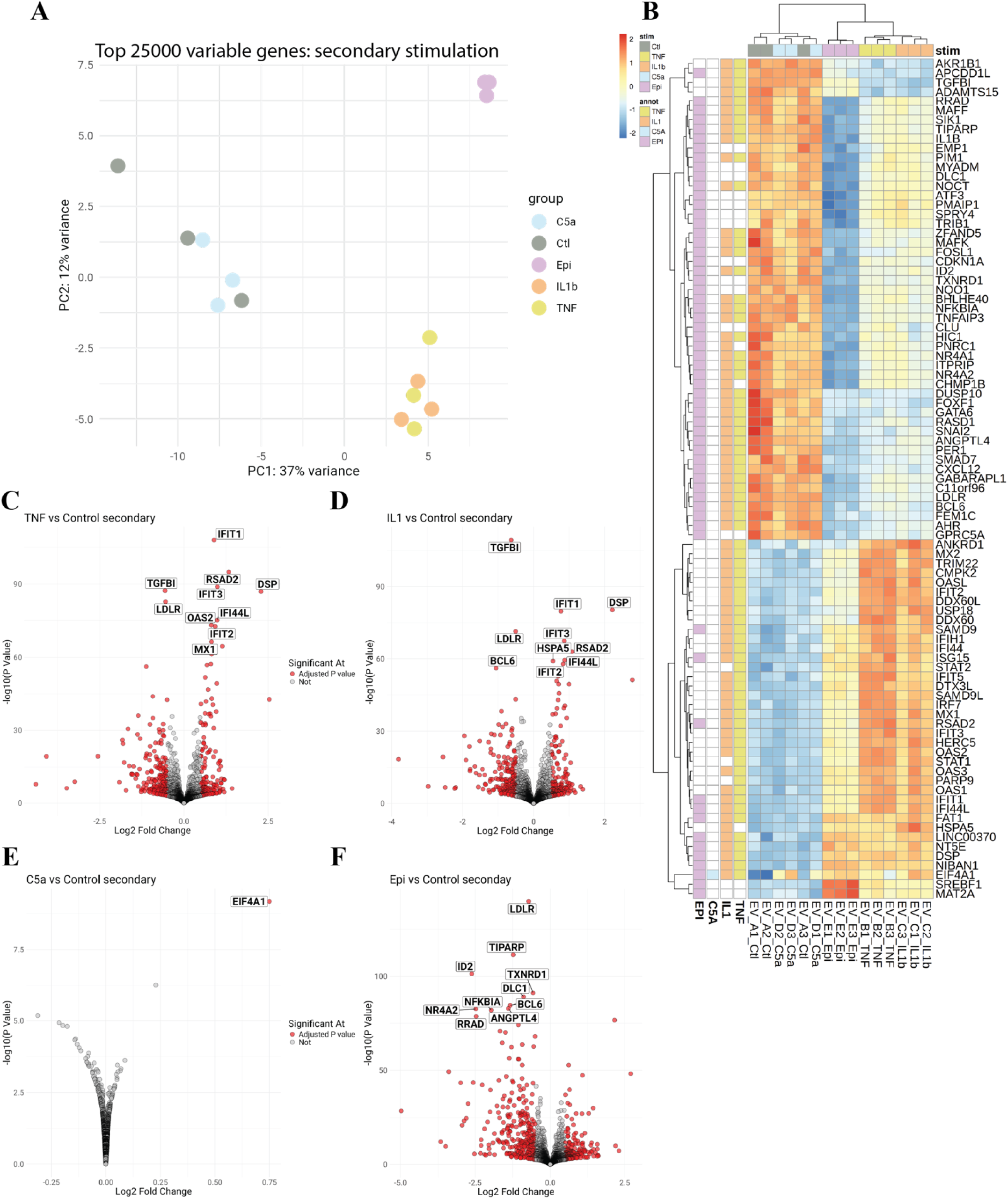
Secondary stimulations induced novel patterns of gene expression. A) PCA plot generated with the top 25000 differentially expressed genes among all secondary stimulations (n=3 for each condition). B) Heatmap depicting the top differentially expressed genes among all secondary stimulations, sample annotations are colored by their stimulation and gene annotations are colored if they are significant at all in that stimulation. C-F) Volcano plots of differentially expressed genes from secondary stimulations for C) TNFα, D) IL-1β, E) C5a, and F) epinephrine. The top 10 differentially expressed genes, sorted by adjusted P value, are labeled. Only one gene was significantly differentially expressed in C5a secondary stimulations.

### 3.4 Overlapping Transcripts of Primary and Secondary Stimulations

One of the key questions we sought to answer was whether any aspect of the transcriptomic response following primary stimulation is propagated by EVs in a feed-forward, autocrine fashion. Therefore, we analyzed overlapping DEGs between primary and secondary stimulations, looking for genes which are differentially expressed in the same direction in both primary and secondary stimulations. C5a stimulation did not significantly differ from no stimulation; the only DEG following secondary C5a stimulation which met criteria for inclusion was *EIF4A1*. TNFα stimulation, both primary and secondary, induced five overlapping genes which were either induced by or inducers of IFN signaling, *ZCH3HAV1, IFIT2, GBP1, IFI16,* and *APOBEC3G* (Figure S2A). Shared DEGs in TNFα primary and secondary stimulations were also related to inflammation and endothelial cell activation, namely, *TNFSF10, TRIM5, LTB,* and *ANKRD1*. Several chemokines are also upregulated in both primary and secondary stimulations in the TNFα and IL-1β groups, namely, *CXCL5, CXCL6, CXCL10, CXCL11,* and *CX3CL1*. Of note, *CXCL10* and *CXCL11* encode IFN-γ–inducible chemokines^18^.

Interestingly, epinephrine primary and secondary stimulations resulted in differential expression of several channel and transporter genes, *SLC6A15, SLC12A8, SCN9A, SLC25A25,* and the antisense RNA *SLC30A4-AS1*; these are mostly solute carrier family ion transporters. When the sets of upregulated and downregulated genes are plotted on Venn diagrams displaying all DEGs in each condition, the extent of overlap is more clearly visualized (Figure S3A-H). C5a, in contrast, did not have any DEGs which overlapped, consistent with the small transcriptomic response to secondary stimulation.

## 4 Discussion

The work reported here represents a first step in understanding how soluble factors affect BBB integrity and in developing EV-based readouts of BBB health. We have identified DEPs in BMVEC-EVs which are common to all EVs produced after BBB-breaching stimuli and may serve as the basis for a serum EV-based readout of BBB integrity. While neuroimaging techniques have advanced such that gadolinium is no longer required to assess BBB permeability, even non-invasive modalities such as a combination of arterial spin labeling and diffusion-weighted imaging are still time-consuming, costly, and not universally available^19–21^. This work suggests that one could use serum-based BMVEC-EVs to assess the functional health of the BBB. Further development in this regard relies on maturation of technologies for non-destructive and specific isolation of BMVEC-derived EVs from serum as well as validation of the markers identified in this study.

EV protein cargoes following the primary stimulations in our study are distinct, suggesting stimulus-specific control of EV cargo loading. There are clear differences between clustering patterns when comparing transcriptional profile of stimulated BMVECs and the protein cargo of their EVs; IL-1β is more similar to C5a in resultant EV protein composition than it is to TNFα, despite similarities in transcriptional responses between IL-1β and TNFα.

Furthermore, there is variance in the number of unique proteins in these EVs depending on stimulus. These data suggest differential, stimulus-specific selection of the proteins which are packaged into EVs and exported from cells following BMVEC activation, with a minority of upregulated genes contributing to the EV protein cargoes. Thus, a diversity in protein cargoes exists despite similarities in transcriptomic responses. Most importantly, the fact that some proteins are expressed in EVs from BMVECs after all stimulations but not EVs from unstimulated BMVECs strongly suggests potential diagnostic utility Although we demonstrate how different stimuli result in unique protein cargoes in associated BMVEC-EVs, we do not provide a mechanistic explanation for this differential cargo loading. Several other groups have described how inflammation alters the characteristics and function of EVs, with Wang et al. demonstrating a link between TNFα and glutaminase in regulating the release of EVs from astrocytes in the context of inflammation^22^. Differential regulation of lipids may also contribute to alterations in EV cargo selection; the presence of lipid raft-associated proteins and molecules, alongside blockage of secretion of specific proteins such as αB-crystallin upon disruption of lipid rafts, implicates these membrane components as drivers of EV cargo loading^23^.

We identified several functional motifs in EV-associated proteins common to several or all stimulations. Downregulation of proteins related to tight junctions and BBB structural integrity is shared among all primary stimulation responses, consistent with a large body of literature which reports that inflammation decreases tight junction protein expression. We identified S100A11 in TNFα-stimulated EVs which has known functional significance when EV-associated^24^. S100A11 is downstream of NF-κB and it has immunomodulatory functions when secreted in EVs, stimulating release of chemokines from peripheral immune cells^24–26^. RALB, also in EVs of TNFα-stimulated BMVECs, induces endothelial cell activation and subsequent release of von Willebrand Factor^27^.

We identified a set of differentially expressed genes which were common to all the BBB-breaching stimuli we studied and within the top 100 DEGs of at least one stimulus. These common DEGs include those encoding tight junction proteins and tight junction-associated proteins, which were downregulated in all conditions, as well as genes encoding several transport proteins. TNFα and IL-1β both engage pathways which may drive direct disruption and inflammatory activity at the BBB. C5a also activates NF-kB, like TNFα and IL-1β, but is distinct in its enrichment for zinc finger proteins. Epinephrine is unique in its induction of genes encoding regulators of vascular tone, angiogenesis, and transport channels. A number of functional categories and specific transcripts were represented in both primary and secondary responses.

The substantial presence of chemokines and related genes in both the primary and secondary responses of BMVECs to TNFα and IL-1β suggest that lymphocyte penetration of the BBB may be a consequence of exposure to elevated concentrations of these cytokines^28–32^. Interestingly, the extracellular matrix of the BBB sequesters chemokines to increase their local concentration^33^. TNFα and IL-1β upregulate expression of various MMPs and acute-phase proteins which are implicated in the breakdown of tight junction proteins in the context of inflammation, infection, and stroke^34–36^. TNFα DEGs in both primary and secondary signatures include genes enriched in type 1 IFN and IFN-γ signaling pathways. The presence of IFN-related signatures in TNFα and IL-1β responses provide further suggest that IFNs and their signaling pathways are important regulators of BBB function in inflammatory states. IFN-γ has been described to bolster the BBB and has also been described to disrupt BBB function^37,38^.

C5a disrupts the BBB through binding to CD88 and inducing NF-kB signaling and subsequent production of reactive oxygen species, transcription of iNOS mRNA, and cytoskeletal remodeling^39–43^. Inhibition of C5a ameliorates BBB dysfunction in the setting of sepsis^44^. We found that the primary response to C5a included upregulation of several zinc finger proteins, which have been described to enhance BBB function in the context of stroke and increase expression of tight junction proteins such as occludin^45^. The downstream effectors of one of these zinc fingers, KLF2, are involved in attenuating inflammation-induced endothelial adhesion molecules VCAM-1 and E-selectin in addition to inducing endothelial nitric oxide synthase expression, which regulates vascular tone and cerebral blood flow^46^. Likewise, KLF11 reduces endothelial cell activation, increases tight junction expression, and decreases expression of cellular adhesion molecules downstream of NF-κB-p65^47,48^. Zinc finger proteins are also implicated in protection of the CNS against oxidative stress in ischemic stroke and hypoxia^49^.

The upregulation of zinc-finger proteins suggests the existence of a BBB-protective transcriptomic profile following C5a stimulation. The responses of BMVECs to EVs from C5a and unstimulated EVs do not differ significantly in spite of the 313 detected proteins which are distinct between these EV populations, providing evidence that EV-associated proteins are unlikely to be crucial in feed-forward signaling after exposure of BMVECs to C5a.

Epinephrine has been proposed to disrupt BBB integrity indirectly through alterations in cerebral hemodynamics; the BBB of the amygdala is selectively permeable in NPSLE and epinephrine could be a driver of this permeability^50–52^. We found that epinephrine not only induced genes related to nitric oxide and smooth muscle regulation of vascular tone but also induced expression of several genes encoding transmembrane transporters and channels in both primary and secondary stimulations. Several angiogenesis-related factors were also expressed in both primary and secondary epinephrine stimulations. This may suggest that epinephrine-induced BBB dysfunction is not solely mediated through epinephrine’s effects on vascular hemodynamics but also by the direct action of epinephrine on BMVECs through other pathways^53^.

Although a BMVEC monoculture allows for certainty in the source of EVs, it also lacks intercellular interactions present in the neurovascular unit, namely the contributions of pericytes and astrocytes, and does not experience the same mechanical stresses as the in-vivo BBB^54^. A more robust BBB model, such as with simulated blood flow, is a future need.

## Author Contributions

**Alec Z. Xiang:** conceptualization, methodology, software, validation, formal analysis, investigation, data curation, writing - original draft, writing - review and editing, visualization, project administration. **Zara Adeel:** methodology, investigation, validation, visualization, writing - review and editing. **Andrew J. Shih:** conceptualization, methodology, software, validation, formal analysis, data curation, writing - original draft, writing - review and editing, visualization. **Betty Diamond:** conceptualization, methodology, resources, writing - original draft, writing - review and editing, supervision, project administration, funding acquisition.

## Acknowledgements

We thank the lab of Joan W. Berman for their generous gift of primary human astrocytes and their technical expertise in coculture systems. Transmission electron microscopy images were acquired by the Analytical Imaging Facility at the Albert Einstein College of Medicine and RNA sequencing data was generated by Novogene. Mass spectrometry data was generated by the Proteomics Resource Center at The Rockefeller University with specific thanks to Henrik Molina for discussion and technical assistance. Portions of the presented research was conducted as part of the doctorate dissertation work of the first author at the Donald and Barbara Zucker School of Medicine at Hofstra/Northwell.

## Data Availability Statement

The authors confirm that the data supporting the findings are available within the article, its supplementary material, or deposited to the GEO (GSE315626) or PRIDE (PXD072771). Other data that support the findings reported in this study are available from the corresponding author upon reasonable request.

