## Supplemental Materials for "Blood-brain barrier disrupting stimuli induce production of extracellular vesicles with distinct protein cargoes and functionality"

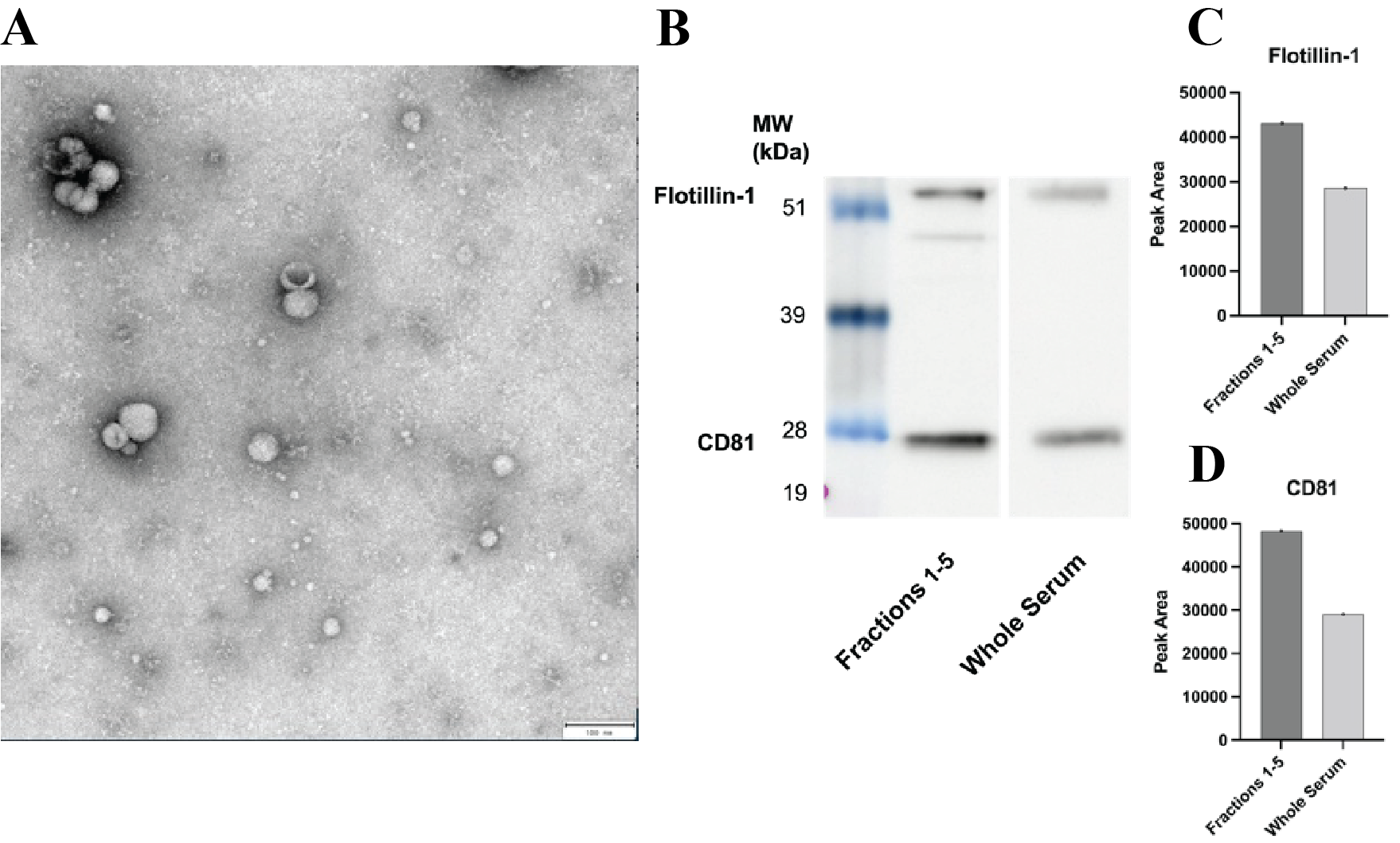

**Figure S1. EVs are enriched from cell culture supernatant.** Aliquots of cell and organelle-depleted BMVEC cell culture supernatant were processed to obtain EVs. A) Transmission electron microscopy showing vesicles which exhibited classical morphological features of EVs, including the presence of cup-shaped disks and diameters typical of EVs. EVs from cell culture supernatant were imaged, and all samples contained abundant EVs; a representative image exemplifying this characteristic morphology is shown. B) Enrichment of EVs was confirmed by performing a Western blot for flotillin-1 and CD81, comparing signal from human serum-derived EVs against cell-depleted human serum. C) Densitometric quantification of Flotillin-1 signal. D) Quantification of CD81 signal.

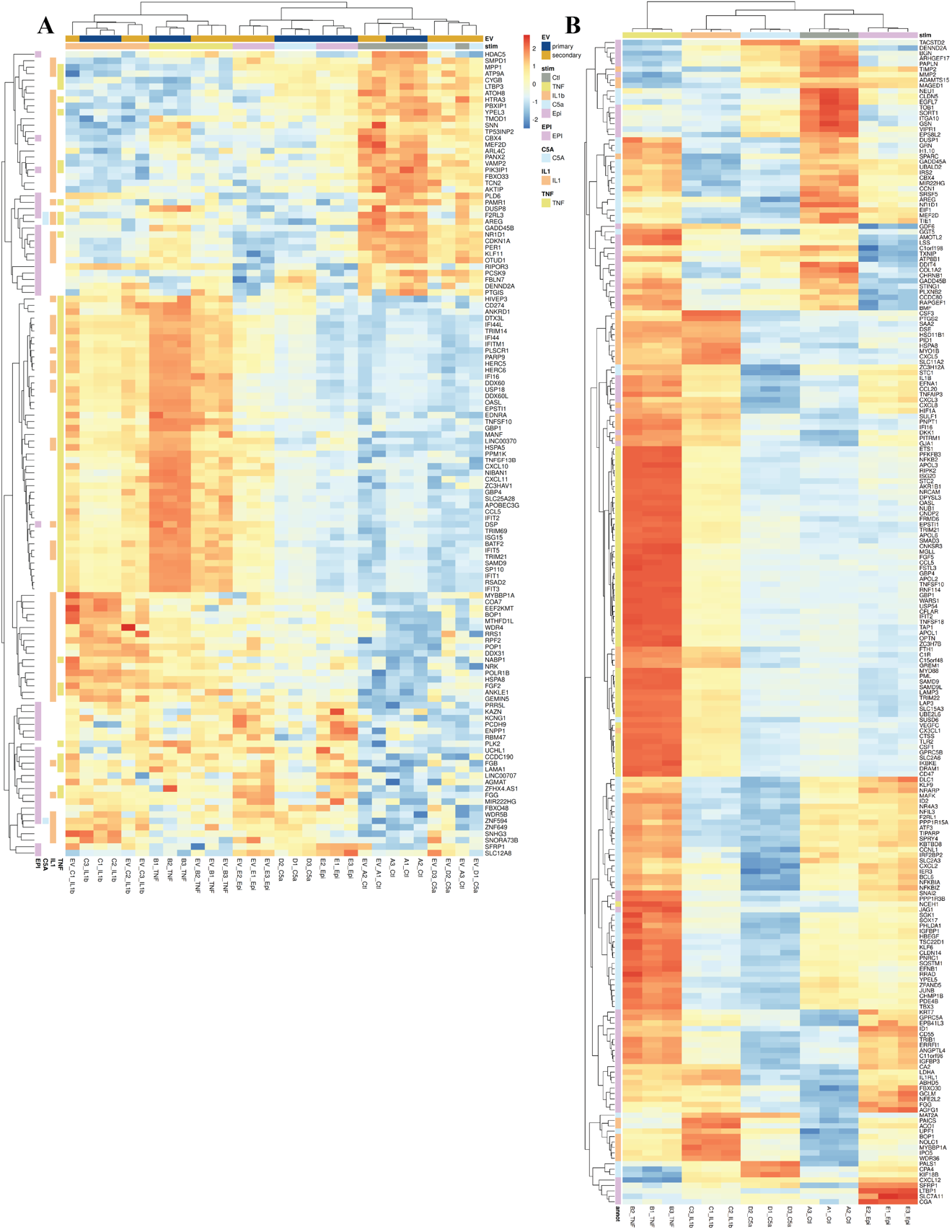

**Figure S2.** **Visualizing overlapping and fingerprint DEGs.** A) Heatmap displaying genes which are differentially expressed in both primary and secondary stimulations for each stimulus. B) Heatmap displaying the genes unique for each condition within the top 100 differentially expressed genes of each specific condition.

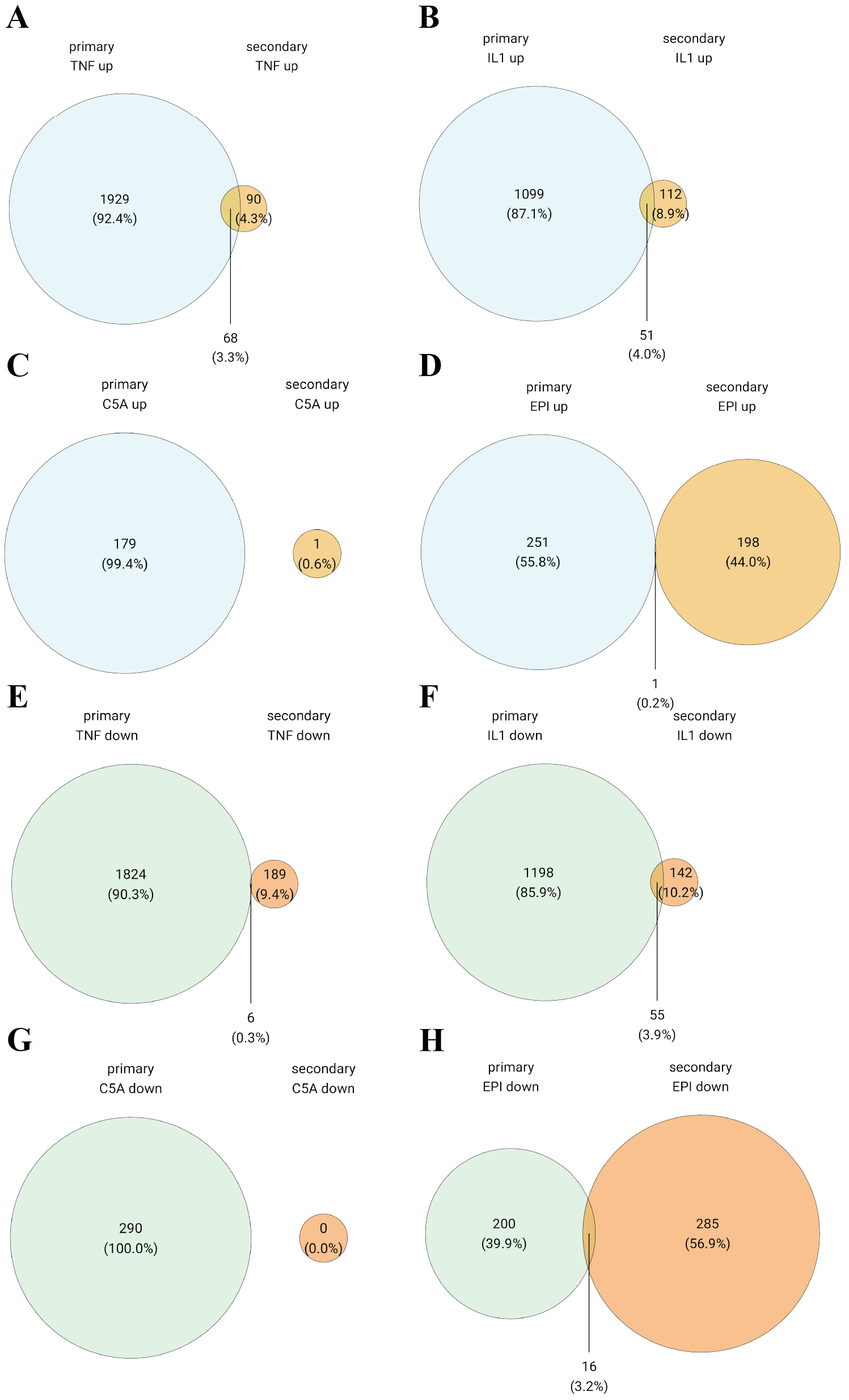

**Figure S3. Primary and secondary stimulations induced expression of distinct but sometimes overlapping sets of genes.** A-D) Venn diagrams of upregulated genes in A) TNFα, B) IL-1β, C) C5a, and D) epinephrine primary and secondary stimulations with the overlapping region representing the number of genes common to both stimulations. E-H) Venn diagrams of downregulated genes in E) TNFα, F) IL-1β, G) C5a, and H) epinephrine primary and secondary stimulations with the overlap representing the number of genes common to both stimulations.

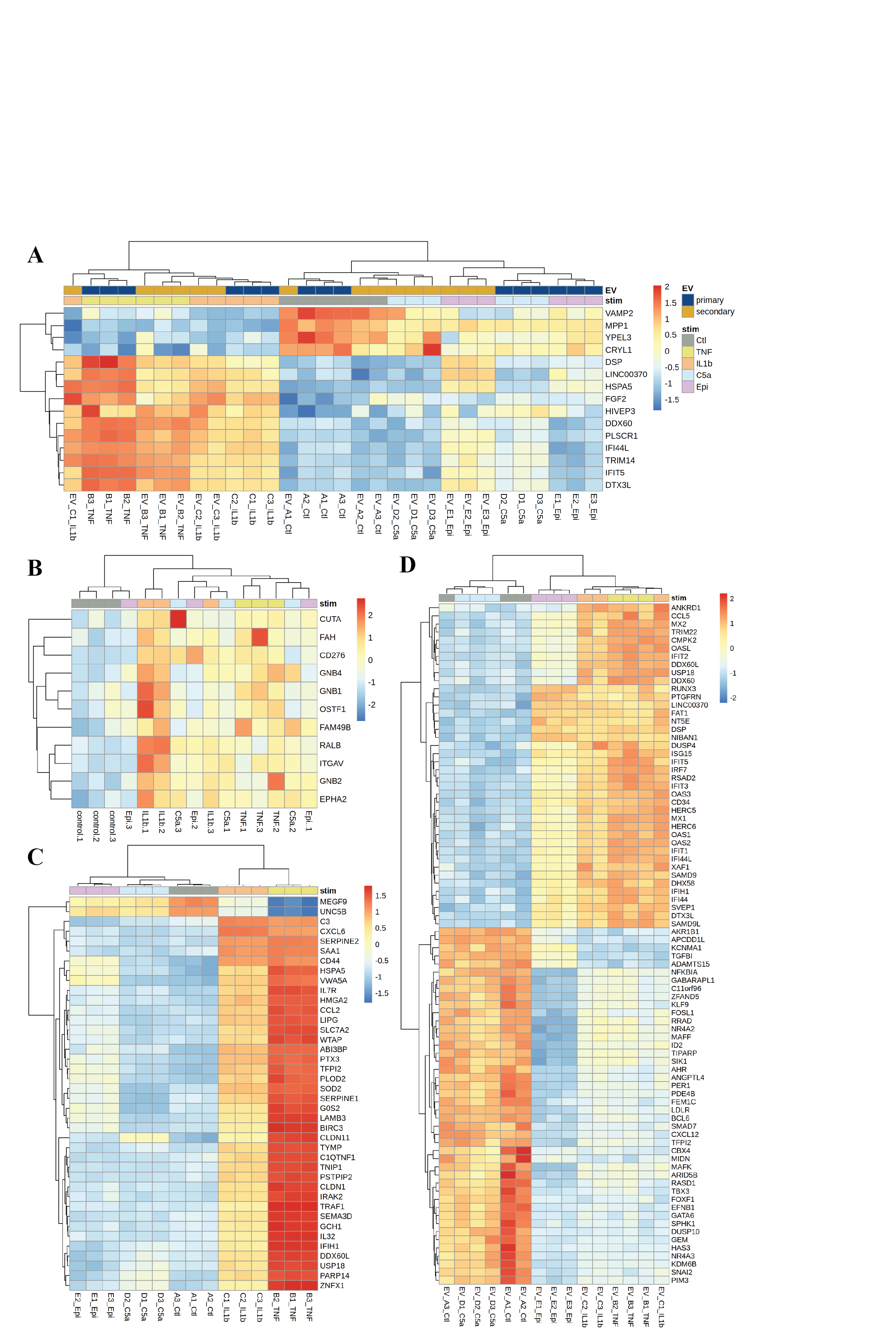

**Figure S4. Mapping overlap between TNFα and IL-1β** A) Heatmap of top 100 DEGs which are expressed in both primary and secondary stimulations of TNFα and IL-1β**.** B) Heatmap of DEPs shared between EVs resulting from TNFα and IL-1β stimulations. C) Heatmap of DEGs which are common between TNFα and IL-1β in primary stimulations. D) Heatmap of DEGs which are common between TNFα and IL-1β in secondary stimulations.

**Table S1 – Common DEGs in Primary Stimulations**

| feature | TNFα | IL-1b | C5a | Epinephrine |
| --- | --- | --- | --- | --- |
| TGFB2 | 1.399 | 1.190 | 0.554 | 0.625 |
| TCN2 | -0.731 | -1.100 | -0.664 | -0.684 |
| AREG | -1.075 | -0.982 | -2.610 | -0.899 |
| GDF15 | -0.886 | -0.898 | -0.967 | -0.725 |
| RGS4 | 0.814 | 0.732 | 0.902 | 0.996 |
| CLDN5 | -1.241 | -1.501 | -1.273 | -0.862 |
| HDAC5 | -0.656 | -0.894 | -0.863 | -1.250 |
| NR1D1 | -1.004 | -1.372 | -1.984 | -1.046 |
| EGFL7 | -1.098 | -0.975 | -0.812 | -0.651 |
| NEU1 | -0.509 | -0.677 | -0.666 | -0.519 |
| FAM43A | -0.830 | -0.803 | -0.794 | -1.260 |
| FBXO30 | 0.924 | 0.781 | 0.520 | 1.035 |
| SFRP1 | 0.638 | 1.191 | 1.757 | 2.464 |
| MMP2 | -2.417 | -1.467 | -0.773 | -0.991 |
| VIPR1 | -2.448 | -2.341 | -1.298 | -1.742 |
| TOB1 | -0.560 | -0.866 | -0.532 | -0.797 |
| DKK1 | 1.599 | 1.107 | 0.635 | 0.756 |

A table of genes which were differentially expressed across all primary stimulations and within the top 100 DEGs of at least one stimulation. Values represent log 2 fold changes.
